# Membrane localization accelerates association under conditions relevant to cellular signaling

**DOI:** 10.1101/2022.10.22.513358

**Authors:** William Y. C. Huang, Steven G. Boxer, James E. Ferrell

**Author notes:** Corresponding author. (W.Y.C.H.), (J.E.F.).

## Abstract

Translocation of cytoplasmic molecules to the plasma membrane is commonplace in cell signaling. Membrane localization has been hypothesized to increase intermolecular association rates; however, it has also been argued that association should be faster in the cytosol because membrane diffusion is slow. Here we directly compare an identical association reaction in solution and on supported membranes. The measured rate constants show that for 10 μm-radius spherical cell, association is 15-25-fold faster at the membrane than in the cytoplasm. The advantage is cell size-dependent, and for typical ~1 μm prokaryotic cells it should be essentially negligible. Rate enhancement is attributable to a combination of closer proximity of the signaling molecule to its targets after translocation and the higher efficiency of a two-dimensional search.

## Main Text

The translocation of signaling molecules from the cytosol to the cell membrane represents a key step in many signaling pathways. The process is exemplified by Ras activation in receptor tyrosine kinase (RTK) signaling (Fig. 1A) (*1*–*3*). Autophosphorylated transmembrane receptors recruit the nucleotide exchange factor Son of Sevenless (Sos) from the cytosol via one (Grb2) or two (Shc plus Grb2) adaptor proteins. Membrane-associated Sos then activates membrane-bound Ras protein, which in turn activates the MAP kinase (MAPK) cascade and initiates transcriptional programs for proliferation, differentiation, cell motility, and/or survival. Despite the fact that Sos activation is a complex, multistep process (*4*), the artificial recruitment of Sos to the membrane in receptor-independent manner is sufficient to activate the MAPK pathway and downstream transcriptional reporters (*5*, *6*). A wide variety of other signaling proteins also translocate to the membrane to become activated and/or to find targets, including phospholipase C gamma (PLCγ), phosphatidylinositol 3-kinase (PI3K), and both 3-phosphoinositide-dependent protein kinase 1 (PDK1) and its target Akt (*7*, *8*). Thus, membrane translocation is a recurring theme in eukaryotic signaling.

**Fig. 1.**
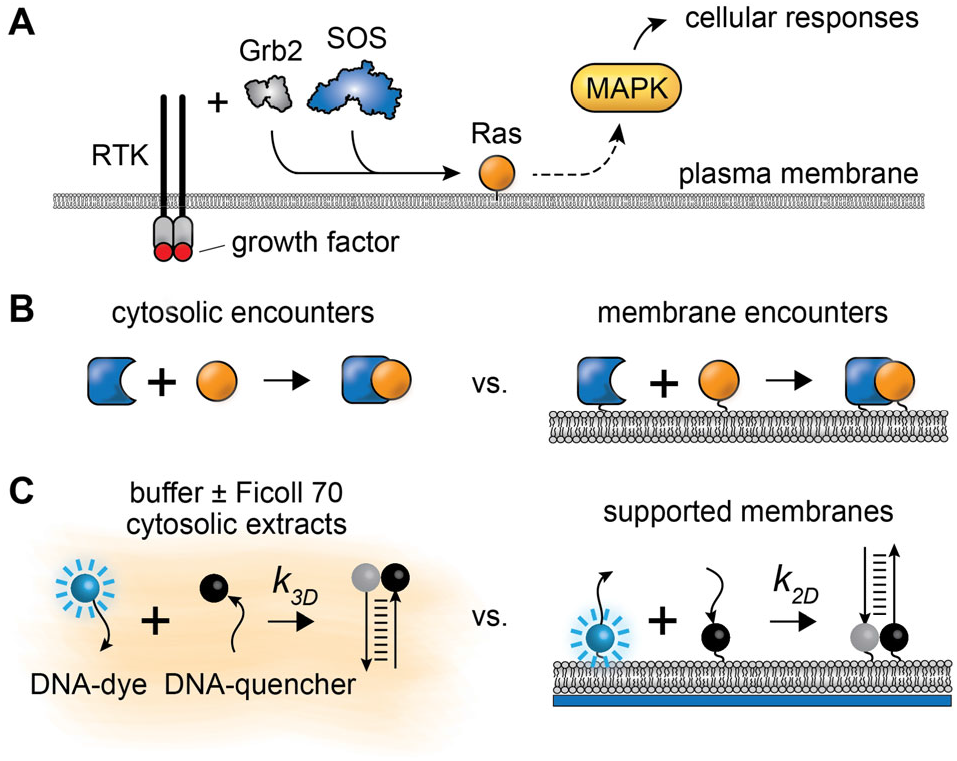
Molecular association in 2D vs. 3D in cellular signaling. (**A**) Simplified schematic of a receptor tyrosine kinase (RTK) signaling pathway. The growth factor brings about dimerization, activation, and intracellular autophosphorylation of the RTK, which recruits cytosolic proteins to the membrane. Sos, a key GEF protein for Ras GTPase activation, is recruited to the cell membrane via the adaptor protein Grb2. Membrane-associated Sos activates Ras, which activates the MAP kinase cascade. (**B**) Schematic of association in 3D in the cytosol vs. 2D on a membrane. (**C**) Our approach to directly compare rates in 2D vs. 3D: a controllable DNA association reaction monitored on supported membranes, in buffers with or without viscogens, and in cytosolic extracts.

This raises the fundamental question of how membrane localization promotes signaling. The potential advantages of a two-dimensional (2D) search over a three-dimensional (3D) search have long been of interest to theoreticians (*9*–*15*). For example, Pólya’s theorem shows that random walks on a grid in 2D will always find a stationary target, but in 3D the probability is less than one (*9*). Adam and Delbrück further explored the question and hypothesized that translocation of molecules from the cytoplasm to a cell membrane increases first-encounter rates and association rates (*10*). This concept has been elaborated on in a large number of subsequent papers (*11*–*14*, *16*–*18*). In the framework of Michaelis–Menten kinetics, a faster association rate could directly speed the activation of the target if the process is diffusion-controlled, or increase the steady-state concentration of protein-target complexes and thereby increase the rate of target activation if the process is reaction-controlled.

However, membrane translocation would also be expected to slow diffusion because membranes are more viscous than cytoplasm. Whether or not this slowing would completely counteract the local concentration effect is currently a matter of conjecture (Fig. 1B), with some estimating that a membrane-associated 2D search should be quicker than a cytoplasmic 3D search (*10*, *14*, *17*),and others predicting the opposite (*16*, *19*).

Here we set out to resolve these uncertainties through experiments comparing an identical association reaction in solution and at the membrane. We chose a model system approach to enable quantitation under controlled conditions: buffer solutions and cytosolic extracts to mimic the physical environment of the cytoplasm, and supported phospholipid bilayers to represent cell membranes (Fig. 1C). Each of these systems has a long history of generating mechanistic insights into biological processes occurring in cellular environments (*3*, *20*, *21*). Supported membranes preserve the hallmark lateral fluidity of cell membranes, with diffusion largely determined by the viscosity of the lipids. Cytosolic extracts retain the composition and concentrations of cytosolic molecules with essentially no dilution. Dynamics measured in extracts can also serve as a benchmark for experiments in simple buffer solutions containing crowding agents or viscogens. The association reaction we chose to study was the binding of protein-sized complementary DNA strands to each other (Fig. 1C). By coupling a fluorescent dye to one strand and a fluorescent quencher to its complement, association can be monitored as fluorescence decay in real time (Fig. 1C and S1A-C) (*22*). Because short DNAs are relatively predictable in terms of structure, thermodynamics, and kinetics (*22*–*24*), they are particularly useful for elucidating the dynamical effects of membrane localization. DNA binding reactions are also robust and relatively easy to control. In addition, the slow dissociation rates of double-stranded DNA allow association rates to be determined directly, simplifying the analysis. In the following experiments, we monitored association reactions over timescales of ≲ 10^3^ s, whereas the average lifetime of the DNA complexes used was ≳ 10^6^ s (Fig. S1D).

The use of DNA enables us to initiate the binding reaction through strand-displacement reactions (*22*–*24*). Measuring association rates on membranes is potentially problematic because the DNA strands could associate during the coupling of the fluorescent (F) and quencher (Q) strands to membrane lipids. To prevent this, strands were protected with complementary DNAs: a blocker strand (B) for the fluorescent strand, and a quencher anchor (A) for the quencher strand (Fig. 2A), using DNA sequences adapted from previous work on lipid-lipid encounters (*22*) (Fig. S1A). The F and A strands were functionalized with sulfhydryl groups to allow them to be covalently coupled to maleimide-derivatized lipids (*25*). The association reaction of interest, F to Q, was then initiated by addition of an initiator strand (I) with a higher affinity for the blocker (B) than the blocker had for the fluorescent strand (F). The initiator binds to the toehold region on B and then outcompetes F in a strand displacement process. F is then free to diffuse and bind to its complementary strand Q through a second toehold-mediated strand displacement process. Blocking was found to protect about 80% of strands and the protection remained stable over hours. Although strand-displacement reactions are not necessary to time-resolve reactions in solutions, we followed this identical protocol for consistency.

**Fig. 2.**
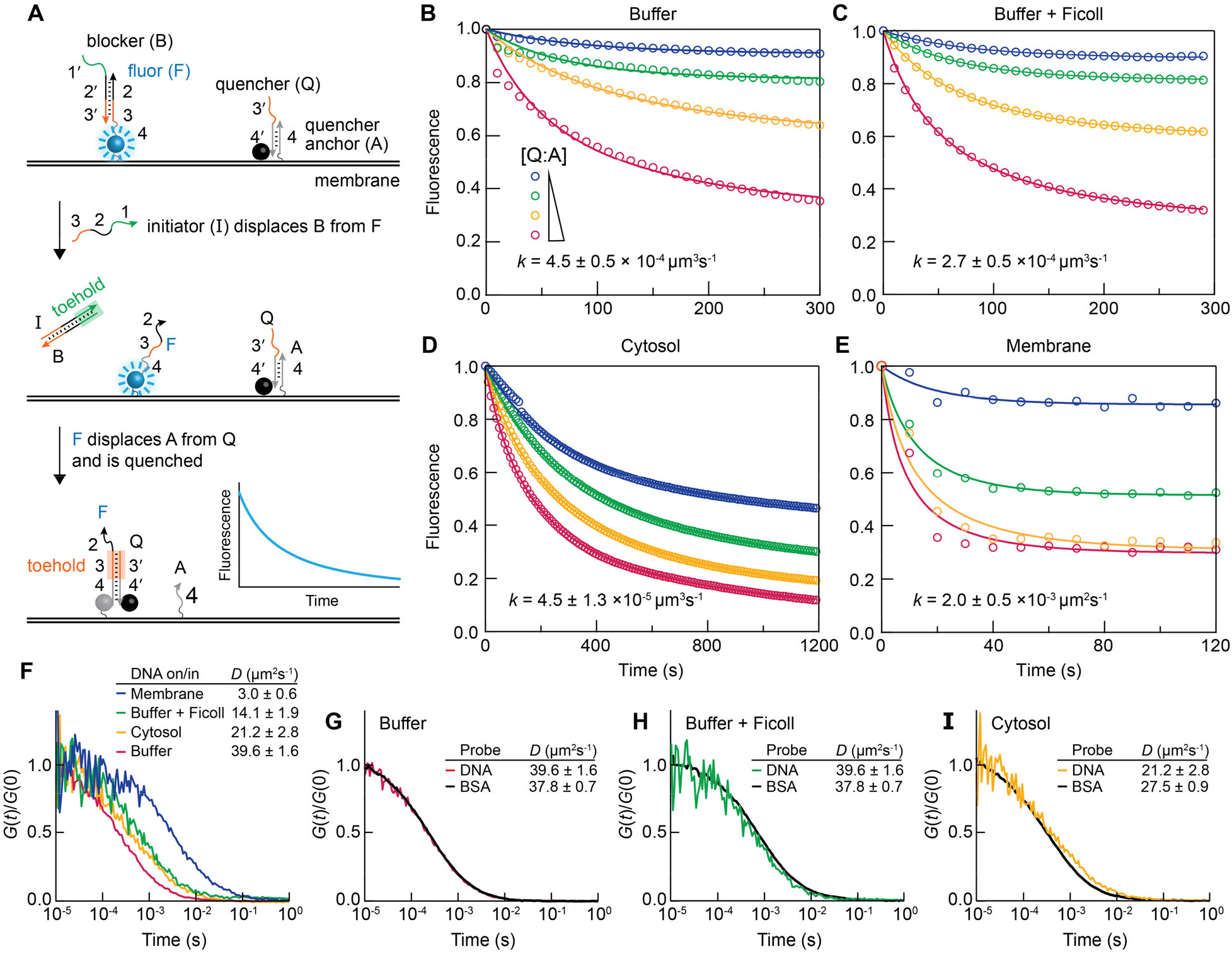
Association reactions and diffusion measurements. (**A**) Schematic view of the strand displacement reaction. (**B-E**) Normalized traces of fluorescence as a function of time after initiating the of DNA strand-displacement reactions in phosphate buffered saline (**B**), phosphate-buffered saline plus 10% (w/w) Ficoll 70 (**C**), cytosolic *Xenopus* egg extracts (**D**), and on supported membranes (**F**). In these titration experiments, the fluorescent complex (B:F) was fixed to about 50 nM, 100 nM, 250 nM, and 50 molecules·μm^-2^ for (**B**-**E**), respectively. The quencher complex (Q:A) in solution titration was {50, 25, 10, 5}, {100, 50, 25, 10}, and {250, 200, 150, 100} nM for (**B**-**D**), respectively; for membrane experiments, the quencher was incubated at {1x, 0.75x, 0.5x, 0.25x} of the fluorescent complex during the coupling reaction. Membrane data shown here have been corrected for photobleaching (Fig. S5). Solid curves are fits of the equation 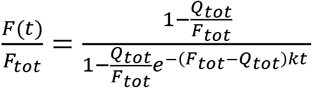, where *F_tot_* is the total concentration or surface density of the fluorescent DNA strand, *Q_tot_* is the total concentration of the quencher, and *k* is the association rate constant. The rate constants are averages ± S.E.M. with *n* = 4. (**F**) Normalized FCS autocorrelation functions of DNA F:B complexes. Diffusion coefficients were obtained from fitting a 3D Brownian model to the data, except in the case of membranes, in which a 2D Brownian model was used. Diffusion coefficients are shown as fitted values ± 95% confidence intervals fitting. (**G-I**) Normalized FCS autocorrelation functions for DNA F:B complexes to BSA-Alexa Flour 488 (black) in buffer (G), buffer plus 10% Ficoll 70 (H), and cytosolic *Xenopus* egg extracts.

Using this approach, we determined the rate constants for association in solution and at the membrane by titration experiments (Fig. 2B-E). For the solution reactions (Fig. 2B-D and S1), fluorescence was measured in a fluorometer; for the membrane experiments (Fig. 2E) we used epifluorescence microscopy. To generate a family of kinetic traces, we varied the concentration of the quencher complex [Q:A] between 0 and 100% of the fixed total concentration of [B:F]. As a control, we measured the photobleaching rate in the absence of the quencher complex (Q:A), which accounted for less than 5% of total molecules (Fig. S1F); thus, most of the fluorescent decay was due to DNA binding. Because the reaction consists of two sequential strand-displacement reactions (I displacing B and F displacing Q), we simplified the reaction kinetics by conducting these experiments with an excess of the initiator strand (I) as compared with the fluorescent complex (B:F). Under these conditions, the first strand displacement reaction could be regarded as essentially instantaneous, and the rate-liming step was the binding of the fluorescent strand (F) to the quencher (Q) (Fig. S1E) (*22*). Thus, the overall kinetics could be approximated by a bimolecular reaction with a time course given by:

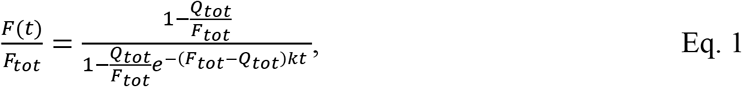

where *F*(*t*) is fluorescence as a function of time with *F*(0) = *F_tot_*, *F_tot_* is the total concentration of the F strand, *Q_tot_* is the total concentration of the Q strand, and *k* is the association rate constant in units of concentration^-1^time^-1^ for 3D solution experiments, and surface density^-1^time^-1^ for 2D membrane experiments. *Q_tot_* and *k* were inferred by nonlinear curve fitting. For supported membrane experiments, *F_tot_* was measured by fluorescence correlation spectroscopy to account for the coupling efficiency (Fig. S2). The solution experiments did not involve covalent coupling reactions, so *F_tot_* was directly known.

For buffer without Ficoll, fitting Eq. 1 to the data yielded a rate constant *k* of 2.7 ± 0.3 × 10^5^ M^-1^s^-1^ (mean ± standard error of the mean (SEM), unless stated otherwise), or 4.5 ± 0.5 × 10^-4^molecules^-1^μm^3^s^-1^ (Fig. 2B). In buffer plus the viscogen Ficoll 70 at a concentration (10% w/w) that yielded diffusion constants similar to those seen in organelle-containing cytoplasm (*26*), titration yielded a rate constant *k* of 1.6 ± 0.3 × 10^5^ M^-1^s^-1^, or 2.7 ± 0.5 × 10^-4^ molecules^-1^μm^3^s^-1^ (Fig. 2C), which is about half of that in buffer without Ficoll. In cytosolic *Xenopus* egg extracts, prepared following standard protocols (Fig. S3A) (*27*), except with the addition of the nuclease inhibitor Mirin and EDTA to minimize probe degradation, titration gave a rate constant of 2.7 ± 0.7 × 10^4^ M^-1^s^-1^, or 4.5 ± 1.3 × 10^-5^ molecules^-1^μm^3^s^-1^ (Fig. 2D), which is about 10-fold slower than buffer. We noticed that in extracts a fraction of the DNA was non-functional in terms of strand displacement reactions, likely due to sequestration by DNA-binding proteins; this non-functional fraction was subtracted from the data before normalization (Fig. S3B). However, we suspect that additional sequestration may be present, perhaps in the form of nonspecific binding with fast exchange, that may lower the free DNA concentration during the course of the measurement (Fig. S3C). Thus, the measured rate is likely lower than the true value. Extract data are discussed further in the *Supplementary Materials and Methods* (Fig. S3).

Next, we measured the association rate on supported membranes (Fig. 2E). The membranes consisted of fluid dioleoylphosphatidylcholine (DOPC) bilayers containing 5% maleimide-derivatized dioleoylphosphatidylethanolamine (DOPE), prepared from sonicated unilamellar vesicles (*Supplementary Materials and Methods*). We then used thiol-maleimide crosslinking to couple DNA strands to the derivatized DOPE and anchor them in the membrane (*25*). We were able to generate a wide range of DNA densities on membranes by this method, from tens to hundreds of molecules per μm^2^, as quantified by fluorescence correlation spectroscopy (FCS) (Fig. S2). After density determination, strand-displacement reactions were initiated and followed by epifluorescence microscopy. Strand-displacement reactions were verified to be functional on supported membranes by checking that the fluorescence decay was dependent on quenchers (Q) and initiators (I) (Fig. S4). For these experiments, photobleaching was non-negligible (~25%), so data were corrected for photobleaching prior to fitting (Fig. S5). We found the reaction to be remarkably fast on membranes (<100 s for ~100 μm^-2^); therefore, most of our experiments focused on lower DNA densities (<100 μm^-2^). Titration with a series of Q:A concentrations yielded a rate constant *k* of 2.0 ± 0.5 × 10^-3^ molecules^-1^μm^2^s^-1^.

We confirmed that DNA molecules (F:B complexes) diffused more slowly on supported membranes than in buffer, buffer plus Ficoll, and cytosolic extracts using FCS (Fig. 2F). In all cases a simple diffusion model fit the FCS autocorrelation data satisfactorily, without the need to invoke anomalous diffusion. The membrane diffusion coefficient (*D*) was 3.0 ± 0.6 μm^2^s^-1^ (95% confidence interval, CI), which is typical of a molecule anchored to a fluid lipid in supported membranes (*28*, *29*). In solution the diffusion coefficients were higher: 39.6 ± 1.6 μm^2^s^-1^ in buffer without Ficoll 70; 14.1 ± 2.1 μm^2^s^-1^ (95% CI) in buffer plus Ficoll; and 21.2 ± 3.1 μm^2^s^-1^ in cytosolic extract. We also compared the diffusion of DNA complexes to the 68 kDa protein bovine serum albumin (BSA) (Fig. 2G-I). In buffer ± Ficoll, diffusion coefficients for DNA and BSA were very similar, suggesting that the DNA construct has a similar length scale (we estimate that the DNA has a long axis of about 11-13 nm; BSA has a long axis of about 14 nm, and based on its partial crystal structure, Sos has a long axis of at least 13 nm, PDB: 3KSY). The diffusion coefficients in 10% Ficoll 70 were close to those for protein probes in cytoplasmic *Xenopus* egg extracts, which contain organelles in addition to cytosolic components (*26*). Diffusion of DNA in cytosol was somewhat slower than expected from the behavior of BSA, again possibly because of binding to cytosolic proteins. Although diffusion in both membranes and solutions was slightly faster than typical values observed in cells (D~1 μm^2^s^-1^ and ~10 μm^2^s^-1^, respectively) (*21*, *30*), the buffer, buffer plus Ficoll, and supported membrane model systems were reasonable approximations of diffusion in cells both in terms of the absolute magnitude of the diffusion coefficients and the fold-differences between solutions and membranes.

Given that diffusion was 5- to 13-fold slower on membranes than in the various solutions, we asked if the association reaction was faster or slower. The rate constants from solutions and membranes cannot be directly compared because of their different units; we therefore calculated total reaction rates. In solution, the reaction rate is given by *k*_3*D*_*C*_1_*C*_2_, where *k*_3*D*_ is the 3D rate constant in molecules^-1^μm^3^s^-1^, and *c*_1_ and *c*_2_ are the concentrations of the fluor and the quencher in molecules·μm^-3^. Units for this reaction rate are molecules·μmV, so the total number of complexes formed per unit time for a hypothetical cell with a volume *V* is *k*_3*D*_*C*_1_*C*_2_*V*. Likewise, the reaction rate at the membrane is *k*_2*D*_*σ*_1_*σ*_2_, where *k*_2*D*_ is the 2D rate constant in molecules^-1^μm^2^s^-1^, *σ*_1_ and *σ*_2_ represent the surface densities of the fluor and quencher in moleculeso μm^-2^. Thus, the reaction rate for a hypothetical cell with a surface area *A* is *k*_2*D*_*σ*_1_*σ*_2_*A*. We then define a dimensionless metric *R*, the rate enhancement factor after translocation to the membrane, as the ratio between total reaction rates at the membrane and in solution:

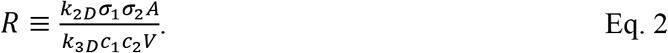

If *R* > 1, association in 2D at the membrane is faster than association in 3D in the cytosol; if *R* < 1, cytosolic association is faster.

If we assume equal total numbers of associating molecules *N*_1_ and *N_2_* in the 2D case and the 3D case (Fig. 3A), then *σ_i_* = *Y_i_*/*A* and *c_i_* = *Y_i_*/*V*. For a roughly spherical hypothetical cell, Eq. 2 becomes:

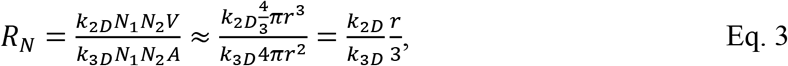

where the subscript *N* in *R_N_* reminds us of the assumption of equal numbers of molecules. Note that the volume-to-area ratio, which determines the degree of condensation from 3D to 2D, is maximal for a sphere; other geometries can be accounted for by using the appropriate volume-to-area ratio (Fig. S6).

**Fig. 3.**
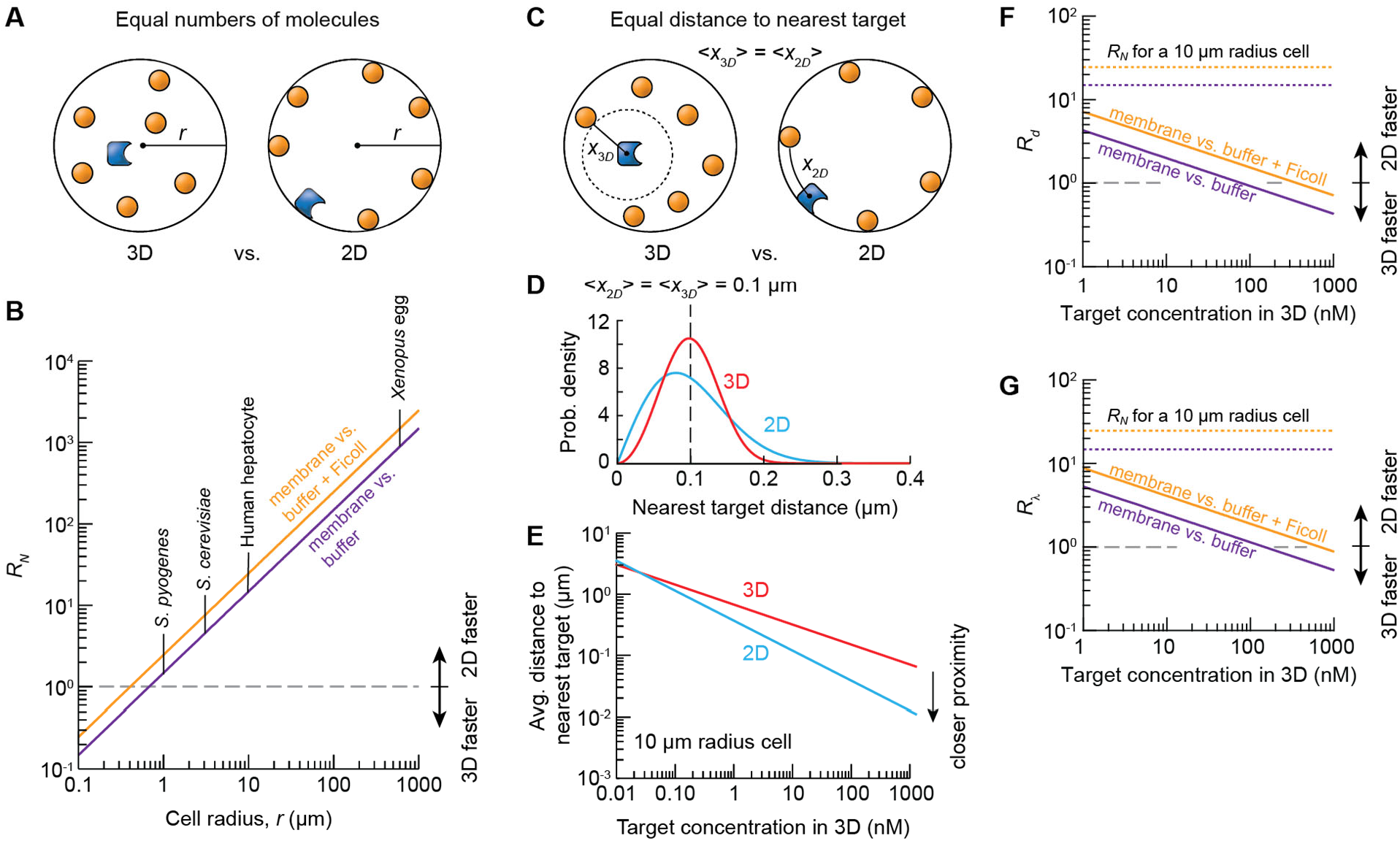
Association reaction rates in 2D vs. 3D. **(A, B)** Changes in association rates when the numbers of molecules are kept constant in 2D and 3D. (**A**) Schematic showing a signaling molecule (blue) and its targets (orange) randomly distributed in the cytoplasm (left) or on the inner aspect of the plasma membrane (right). (**B**) Inferred ratio of the total association rate in 2D divided by the total association rate in 3D (*R_N_*) for spherical cells of various sizes. We assumed equal numbers of molecules in the cytoplasm vs. on the membrane. A value of *R_N_* greater than 1 means that 2D association is faster than 3D association. The diagonal lines are plots of the relationship 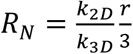 using the value of *k*_2*D*_ from the supported bilayer experiment and the value of *k*_3*D*_ from either the buffer minus Ficoll (purple) or buffer plus Ficoll 70 (orange) data. The radii of one prokaryote and three eukaryotic cells that span a range of sizes are shown. (**C-G**) Changes in association rates from causes other than proximity, calculated by keeping the mean nearest target distance, not the number of targets, the same in 2D and 3D. (**C**) Schematic showing nearest target distances in 2D and 3D. (**D**) Probability density functions for nearest target distances in 2D (red) and 3D (blue), assuming randomly distributed, non-interacting particles. Concentration (for 3D) and surface density (for 2D) values were chosen so that the average nearest target distance would be 0.1 μm for both cases. (**E**) Average distance to the nearest target in a 10 μm radius spherical cell, assuming that the signaling molecule and its targets are cytoplasmic (red) or membrane-localized (blue). (**F**) The ratio of association rates keeping the mean nearest target molecule equal in 2D and 3D, as given by 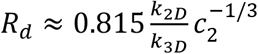. The value of *k*_2*D*_ is from the supported bilayer experiment and the value of *k*_2*D*_ from either the buffer minus Ficoll (purple) or buffer plus Ficoll 70 (orange) data. (**G**) The same as (E) except keeping the linear density (*λ* = c^-1/3^ for 3D and *σ*^-1/2^ for 2D) the same for 2D and 3D.

Eq. 3 shows that the ratio *R_N_* depends upon the size of the cell (Fig. 3B). For a 10 μm-radius spherical cell (4 pL in volume; roughly the volume of a mammalian hepatocyte), association would be 15-times faster if we used the rate constants for membranes (Fig. 2E) vs. buffer without Ficoll (Fig. 2B), and 25-times faster for membranes vs. buffer plus 10% Ficoll (Fig. 2C). If we use the measured rate constants for membranes vs. cytosol, the advantage of 2D becomes higher still (Fig. 2D and S3D), but, as mentioned above, this is probably an overestimate because of the apparent sequestration of DNA strands in cytosol. For a 3 μm-radius budding yeast, *R_N_* is 4 to 8, and for a 600 μm radius *Xenopus* egg, it would be 890 to 1500. For a 1 μm-radius spherical bacterium, there would be little advantage of 2D association over 3D (Fig. 3B). These findings suggest that for most eukaryotic cells, membrane localization substantially promotes intermolecular association, and for many prokaryotic cells it does not.

What accounts for the faster association at the membrane? Here we consider two possible factors (*10*, *14*, *17*). The first is that even though diffusion is slower, the distance between a molecule and its targets has decreased enough to more than compensate, a proximity effect. The second is that the reduction of dimensionality at the membrane makes finding a target easier than it would be in 3D. Either or both of these mechanisms could contribute to the observed facilitation of DNA association by membranes, and their contributions can be estimated through theory.

To determine the extent to which the proximity effect contributes, we begin by calculating the average distance between a signaling molecule and its nearest target molecule in the cytoplasm vs. membrane. We used a statistical approach to derive the probability density *w*(*x*) for random noninteracting particles in 3D (*31*) and 2D (Fig. 3C, D; derivation shown in *Supplementary Methods and Materials*). In 3D, the mean distance 〈*x*_3*D*_〉 ≈ 0.554c^-1/3^, and in 2D, 〈*x*_2*D*_〉 = 0.5*σ*^-1/2^, where *c* is target’s concentration in units of molecules·μm^-3^ and *σ* is in molecules·μm^-2^. These relationships hold irrespective of the size and shape of the cell. For a spherical cell of radius *r*, we can calculate how much the average nearest neighbor distance would change after membrane translocation by using the relationship:

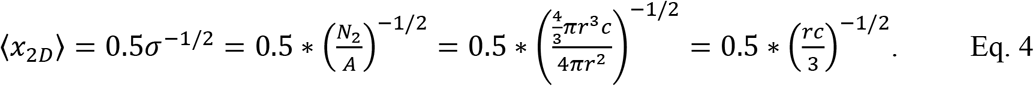

If our spherical cell has a radius of 10 μm, then 〈*x*_2*D*_〉 ≈ 0.274*c*^-1/2^. Note that the mean distance scales differently with concentration in 2D versus 3D, which means that the fold-decrease in distance after translocation is greatest when the target concentration is high (Fig. 3E). Perhaps surprisingly, at very low target concentrations (less than ~10 pM or 24 molecules per cell), the average distance to a target molecule would actually be higher after translocation to the membrane than it had been in the cytoplasm.

The average distance to the nearest target for a target of average concentration (200 nM or 120 molecules-μm^-3^, 1/25,000 of the cell’s total protein concentration (*32*)) would be 112 nm in 3D and 25 nm in 2D, a change of 4.5-fold. Since the time to diffuse a distance *x* is equal to 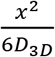 in 3D and 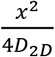 in 2D, concentration alone would be expected to make the 2D association be 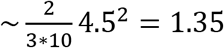-fold faster than 3D association. For a scarcer 10 nM species, the proximity effect would be outweighed by the slower diffusion. However, experimentally, we measured a 15- to 25-fold advantage of 2D over 3D for a hypothetical 10 μm-radius cell, irrespective of the target’s concentration. This indicates that local concentration at the membrane only partly compensates for the slower diffusion, that something other than proximity contributes to the advantage of 2D over 3D, and that it is particularly important for scarce proteins.

To explore this further, we can compare association rates in 2D vs. 3D in the hypothetical 10 μm spherical cell where the mean distance to the target, rather than the number of target molecules, is kept constant (Fig. 3C). The constraint of equal mean distance is satisfied if 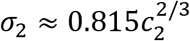. The ratio of the 2D to 3D rates is then:

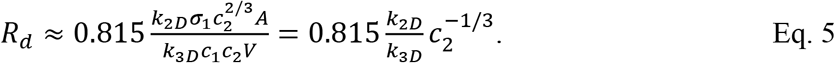

Another way of normalizing for distance is to use equal linear density (*λ*) in 2D and 3D, defined as *λ* = *c*^-1/3^ for 3D and *σ*^-1/2^ for 2D. The ratio of the 2D to 3D association rates at equal linear density is given by:

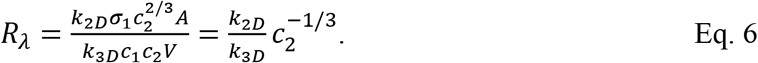

Note that there is less than a 20% difference between *R_d_* and *R_λ_*.

Figs. 3F and G show the distance-normalized metrics *R_d_* and *R_λ_* as a function of the 3D concentration of the target, given our measured values of *k*_2*D*_ and *k*_3*D*_, using Eq. 5 and 6, respectively. For target concentrations of less than ~100 nM, 2D association is still faster than 3D association even when the mean distance to the nearest target is required to be the same and even though diffusion is an order of magnitude slower. In general, for low concentration targets, association occurs faster if the molecules are localized to the membrane even when the nearest neighbor distances are constrained to be identical; for highly concentrated systems, reactions are faster in solution. Note that many of the proteins involved in membrane translocation, and their membrane-bound targets, are in fact scarce. For example, in several mammalian cell lines, Sos is ~5 nM, Ras is ~130 nM, and Raf-1 is ~20 nM (*33*).

Finally, we ask how this extra advantage arises. Suppose we have a diffusing signaling molecule and target protein of radius *r_T_* at a distance *x* away (Fig. 4A), where we take *x* to be the average nearest target distance. The signaling molecule random-walks in 3D until it first reaches a distance of *x* from its starting position. We then repeat this a large number of times. The fraction of these isotropic random walks that arrive at the target (*F*_3*D*_) will be approximately equal to the crosssectional area of the target divided by the surface area of a sphere of radius *x* centered on its starting point (Fig. 4A): 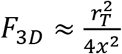. For a 2D random walk, the fraction that hit the target (*F*_2*D*_) will be approximately the diameter of the target divided by the circumference of the circle of radius centered on its starting point: 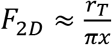. The ratio of these two fractions, 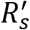 (*s* for search) is:

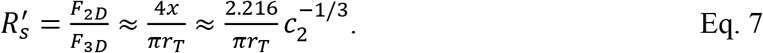

**Fig. 4.**
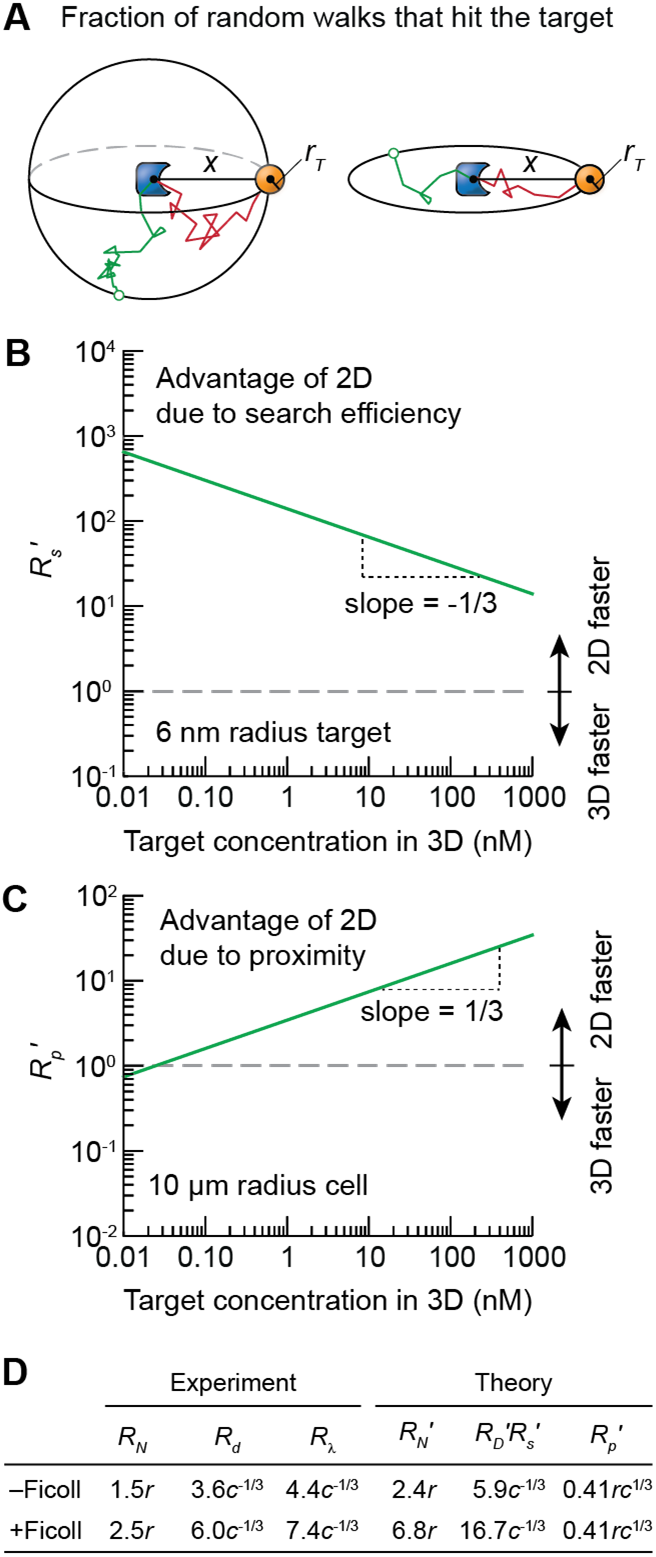
Search efficiency and proximity effects. (**A**) Schematic showing random walk trajectories originating from the center and continuing until they first reach a distance *x*, the mean nearest neighbor distance in 3D (left) and 2D (right). The green trajectories miss the targets; the red trajectories hit them. (**B**) The ratio of the 2D to 3D association rates due solely to changes in the search efficiency as given by 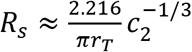. (**C**) The ratio of the 2D to 3D association rates due solely to changes in the proximity of the signaling molecule to its targets after translocation as given by 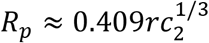. The plot assumed a spherical 10 μm-radius cell. (**D**) Comparison between *R_N_* derived from rate constants and 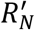 calculated from diffusion 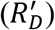, search efficiency 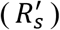, and proximity effects 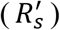. Note that 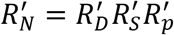. This analysis recapitulates the parameter dependence in *R_N_*, *R_d_* and *R_λ_*.

The prime symbol distinguishes this theoretical rate enhancement factor (*R′*) calculated from fundamental parameters (search efficiency, diffusion, and proximity) from those based on rate constants in Eqs 3, 5, and 6. Note that Eq. 7 accounts for the 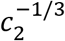 dependence of the extra advantage of 2D association found in Eqs. 5 and 6. 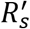 will be greater than one unless the target size is very large and/or the concentration of targets is very high (Eq. 7 and Fig. 4B). For a target protein with a concentration of 200 nM and a radius of 6 nm (half the long-axis of both our DNAs and BSA), only one out of 1,394 3D random walks ending at a distance of 112 nm (the average distance to the nearest target protein) will hit the target. However, in 2D, one out of 59 will, a 24-fold advantage of 2D over 3D. Thus, this calculation predicts that, other things being equal, a 2D search is much more efficient at finding a scarce target than a 3D search is.

To relate search efficiency to the rate constants, we reasoned that a diffusion-dependent association rate is proportional to the fraction of successful random walks divided by a characteristic diffusion time between intermolecular encounters, *F/τ*. The time *τ* is proportional to 〈*x*^2^〉, the mean squared distance over which the diffusion takes place. We then formally define the proximity effect using 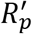 as the fold-advantage of 2D over 3D signaling due solely to the change in the distance between a signaling molecule and its nearest target after translocation (Eq. 4):

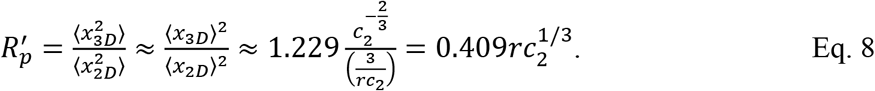

Eq. 8 shows that the proximity effect for a spherical cell depends on both the size of the cell and the concentration of the translocating target (Fig. 4C). Finally, we define 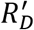 as the fold-change of 2D over 3D due to diffusion rates in 2D vs. 3D. This includes the measured diffusion rates and a factor of 2/3 that arises from the different number of degrees of translational freedom in 2D vs. 3D:

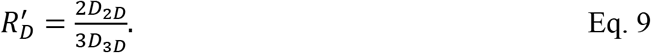

Combining Eq. 7–9 gives

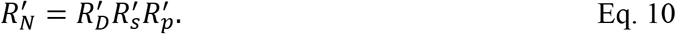

Typically 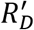 will favor 3D and 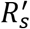 and 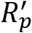 will favor 2D. Note that 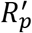 is proportional to the cube root of the target concentration (Eq. 8), whereas 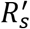 is inversely proportional to it (Eq. 7). As a result, 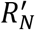 is independent of concentration.

The *R′* values derived from these theoretical arguments agrees fairly well with the experimental values (Fig. 4D), suggesting that diffusion speed, proximity, and search efficiency largely account for the association rates measured in this study. In buffer solution vs. membrane, for example, 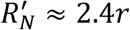 whereas *R_N_* ≈ 1.5*r*. Furthermore, because concentration is accounted for by the proximity effect 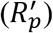, the concentration-normalized metrics, *R_d_* and *R_λ_*, can be compared to 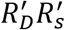. In the same example, 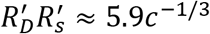 whereas *R_d_* ≈ 3.6*c*^-1/3^ and *R_λ_* ≈ 4.4*c*^-1/3^. Thus, this analysis captures the experimentally-determined enhancement of association by membranes, as well as the relative contributions of proximity vs. search efficiency, to within about a factor of two.

In summary, from measurements of association rate constants for complementary DNA strands whose diffusion dynamics are comparable to those of a typical monomeric protein (BSA), we found that association is generally faster when the strands are anchored to a fluid supported membrane bilayer than when they are free in solution. For a 10 μm spherical cell, the increase in association rate would be 15- to 25-fold, and this would greatly increase the rate of either a diffusion-controlled or a reaction-controlled signaling process. The magnitude of the effect depends upon the size of the cell; for most eukaryotic cells the effect should be substantial, whereas for most prokaryotic cells it should be minimal. Note that regulated translocation to the membrane appears to be commonplace in mammalian cells, and at least one example is known in budding yeast—the translocation of Ste5 between the plasma membrane and the cytoplasm (*34*–*37*). However, we do not know of an example of regulated membrane localization in prokaryotes.

As previously hypothesized, some of the advantage of 2D over 3D can be attributed to a proximity effect. This is particularly true for systems with relatively abundant target proteins. Translocation decreases the mean distance between a protein and its nearest target, and this decrease contributes to an increase in the rate of association. However, much of the advantage of 2D is due to the increase in search efficiency that comes from the reduction of dimensionality: a higher fraction of random walks will hit a target in 2D than in 3D. This is particularly true for scarce target proteins. These findings provide empirical support for the seminal theoretical analysis of Adam and Delbrück (*10*).

Some proteins diffusing in cell membranes may have slower diffusion rates (*D*~0.1 to 1 μm^2^/s) (*30*) than that seen here. But even with a 10-fold reduction in diffusion coefficients, membrane reactions are likely to still be faster than the same reaction in the solution (*R_N_* ≈ 1.5 to 2.5). On the other hand, spatial organization of protein condensates and lipids may increase the effective local concentration of signaling molecules beyond what is produced by simple membrane recruitment, and thereby increase the value of *R_N_* (*38*–*40*). Furthermore, effects other than proximity, search efficiency, and diffusion speed could possibly contribute as well and contribute to the modest difference between the theoretical and experimental results described. For example, the orientations of the associating species could contribute either positively or negatively to the relative advantage of membrane localization: the loss of two presumably unproductive degrees of rotational freedom would favor 2D association, whereas the slower speed of rotation in a viscous membrane environment would favor 3D association. And finally, the viscosity of the membrane, which slows association, would be expected to also slow dissociation, which would increase the steady-state levels of protein-target complexes and thereby promote any reaction-controlled processes. Nevertheless, the present work shows that even in the simple case of essentially irreversible intermolecular association, membrane localization generally has a substantial positive effect on association rates as long as the cell in question is more than one or two microns in radius.

One final question is how the advantage of 2D over 3D would apply to the particular signaling process we started with, the activation of Ras by Sos. If Ras and Sos were both cytoplasmic, there would be only one search involved in the activation of Ras by Sos, a 3D random walk. However, with Ras being constitutively membrane bound, Sos must first find a (scarce) activated RTK and only then perform a 2D random walk to find Ras. A single 3D random walk has been replaced by a 3D random walk plus a 2D random walk. However, Sos is highly processive on membranes: a single activated Sos molecule at the membrane can activate hundreds of Ras molecules before it dissociates (*4*, *41*). Thus, the activation of multiple Ras molecules by membrane-localized Sos instead of cytoplasmic Sos replaces perhaps hundreds of slow 3D searches with one 3D search followed by hundreds of efficient 2D searches. In general, the greater the magnitude amplification generated by a signaling reaction, the greater the advantage of membrane localization.

The present work shows that reduction of dimensionality is implemented differently here than in it is in the interaction of transcription factors with DNA, one of the processes that motivated Adam and Delbrück. In what is now sometimes termed the standard model for the binding of transcription factors to specific enhancer sites on DNA, a combination of 3D searches for a nonspecific DNA strand followed by 1D searches for the enhancer speeds up the target-finding process (*42*). For membrane-associated proteins, processivity means that there are repeated applications of reduction of dimensionality for each of many target searches, thereby amplifying what may otherwise be a weaker 3D-to-2D effect compared to that of 3D-to-1D. Nevertheless, the conceptual similarity between both cases argues that changes in dimensionality, local concentration, and diffusion coefficients may be trade-offs that evolution has weighed and optimized in a number of different types of biological regulation.

## Acknowledgments

We thank Youngbin Lim at the Cell Sciences Imaging Facility for microscopy support, Cheng-ting Tsai for DNA protocols, Xianrui Cheng and Julia Kamenz for extract protocols. We thank John Kuriyan, Jeremy Thorner, and Dan Herschlag for insightful discussions. We thank Yuping Chen, Jo-Hsi Huang, and the rest of the Ferrell lab for helpful discussions and comments on the manuscript.

## Author contributions

W.Y.C.H. and J.E.F. conceived the study. W.Y.C.H. designed, performed, and analyzed the experiments. W.Y.C.H., S.G.B., and J.E.F. interpreted the results. W.Y.C.H. and J.E.F. wrote the paper. All authors commented on the manuscript.

## Data availability

All data is available in the manuscript or the supplementary materials.

## Supplementary Materials

Materials and Methods

Figs. S1 to S6

References (*22*, *25*, *27*, *31*, *43*)

