## Supplementary Materials for "Membrane localization accelerates association under conditions relevant to cellular signaling"

**This PDF file includes:**

Materials and Methods
Figs. S1 to S6
References (*1*-*5*)

### Materials and Methods

DNA Preparation

DNA sequences used in this study are shown in Fig. S1. The sequences mostly followed previous work (*1*), except with the modification of attaching a thiol group for membrane coupling (*2*). All DNAs were synthesized and purified (HPLC) by Integrated DNA Technologies (IDT). Using secondary structure analysis provided by IDT (OligoAnalyzer), we verified that all sequences used have unfavorable hairpin structures ($\Delta G \gtrsim 0$). All DNAs were ordered in single-stranded form. To prepare complexes (B:F and Q:A), we followed protocols suggested by IDT: an equimolar ratio of single-stranded DNAs were mixed and heated to 94°C for 2 min, and then the sample was then let to anneal back to room temperature over an hour. DNA complexes were then stored in a 4°C freezer.

The thiol-containing DNAs were shipped in an oxidized form, protected by an S=S bond. Thus, a reduction step was needed prior to membrane coupling. We followed recommendations by IDT to reduce the thiol group using dithiothreitol (DTT). Before coupling reactions, 5 μM DNA was mixed with 100 μM DTT (Thermo Scientific) in borate buffered saline (BBS) (10 mM sodium borate, 150 mM NaCl, pH 8.5) (Rockland) at 37°C for an hour. The solution was then desalted using Zeba spin desalting columns (7 kDa molecular weight cut-off; Thermo Scientific) three times. Columns were buffer-exchanged to BBS before use. For consistency, we reduced DNA only before membrane coupling reactions in small batches.

Buffer and Crowding Agent Solutions

All experiments (membranes and solutions) were performed in standard phosphate buffered saline (PBS) (11.9 mM phosphate, 137 mM NaCl, 2.7 mM KCl, pH 7.4) (Fisher Scientific), except for cytosolic extracts. To increase buffer viscosity, we added 10% Ficoll 70 (by weight) (Sigma Aldrich) in PBS buffers. Solutions were stirred overnight to ensure proper mixing.

Cytosolic Extracts

Cytosolic extracts were prepared following the protocol of Deming and Kornbluth (*3*), except without the energy regeneration mix. To obtain sufficient cytosolic volume, we typically combined two to three fresh batches of eggs. An additional benefit of combining batches of eggs is that the result may be less specific to one batch of eggs. Because of a higher volume than typical extract preparation, we performed a second round of ultracentrifugation for the cytosolic fraction using the same setting as the first round of ultracentrifugation (250,000 g) for 25 min. Final cytosolic extracts were flash-frozen and kept in a -80°C freezer. All *Xenopus* experiments and animal care followed protocols (APLAC-13307) approved by the Institutional Animal Care and Use Committee (IACUC) of Stanford University. Prior to adding DNA probes, cytosolic extracts were supplemented with 100 μM mirin (Tocris Bioscience) and 20 mM EDTA (Sigma) and incubated for 10 min to inhibit nuclease activities. When adding reagents to solutions (extracts or viscogen agent solutions) for measurements, we minimized dilution by reagents (typically, the solution retained >95% of its concentration, and >90% at the minimum).

Fluorometer

Solution measurements were performed using a JASCO spectrofluorometer (FP-8300 model), which was controlled by the Spectra Manager 2 software. Time course was taken using the Time Course Measurement option in Em Intensity mode. The excitation wavelength was 496 nm (bandwidth 5 nm), and the emission wavelength was 524 nm (bandwidth 5 nm). The response was set to 0.5 s, and the sensitivity was set to medium. Data were typically taken every 10 s. Background fluorescence was always checked prior to measurements. All strand-displacement measurements (solution and membrane) were done at room temperature.

The time-course data were analyzed with a bimolecular reaction: F + Q → F:Q, in which the reaction rate is $kFQ$, where $k$ is the rate constant, $F$ and $Q$ are the concentrations of the fluorescent and quencher strands, respectively. The integrated rate equation for bimolecular reactions has the form

$$\ln\left(\frac{Q/Q_{tot}}{F/F_{tot}}\right) = (Q_{tot} - F_{tot})kt, \qquad \text{Eq. S1}$$

where the subscript *tot* denotes the total (or initial) concentration of $F$ or $Q$. Rearranging Eq. S1 gives

$$\frac{Q}{F} = \frac{Q_{tot}}{F_{tot}} e^{-(F_{tot}-Q_{tot})kt} = \frac{Q_{tot}-(F_{tot}-F)}{F},$$

$$F = \frac{F_{tot}-Q_{tot}}{\left(1-\frac{Q_{tot}}{F_{tot}}e^{-(F_{tot}-Q_{tot})kt}\right)},$$

$$\frac{F}{F_{tot}} = \frac{1-\frac{Q_{tot}}{F_{tot}}}{\left(1-\frac{Q_{tot}}{F_{tot}}e^{-(F_{tot}-Q_{tot})kt}\right)}. \qquad \text{Eq. S2}$$

The last equation of Eq. S2 was used to fit the normalized data. In the fitting procedure, $F_{tot}$ was fixed based on the initial fluorescence intensity, $k$ and $Q_{tot}$ were floating parameters.

Supported Membranes

Small unilamellar vesicles (SUVs) were prepared by mixing 5% (by mol/mol) 18:1PE MCC with DOPC in chloroform, where DOPC stands for 1,2-dioleoyl-sn-glycero-3-phosphocholine and 18:1 PE MCC stands for 1,2-dioleoyl-sn-glycero-3-phosphoethanolamine-N-[4-(p-maleimidomethyl)cyclohexane-carboxamide] (sodium salt) (Avanti Polar Lipids). If visualization of bilayers was required, 0.01% of TR-DHPE was added to the mixture; TR-DHPE stands for Texas Red 1,2-dihexadecanoyl-sn-glycero-3-phosphoethanolamine, triethylammonium salt (Invitrogen). The solution was then evaporated by blowing $N_2$ for 10 min. Dried lipid films were then resuspended in $H_2O$ by vortexing, resulting in a concentration of 0.5 mg/mL. Finally, vesicle solutions were sonicated for 45 s in an ice-water bath twice to make SUVs. SUVs were stored in a 4°C fridge for experiments on the next day. All experiments used new batches of freshly prepared vesicles as described for consistency.

Supported lipid bilayers (SLBs) were assembled in a flow chamber system (μ-Slide, Ibidi). Glass substrates were piranha etched ($H_2SO_4$:$H_2O_2$ = 3:1 by volume; warning: strong acid) (Fisher Scientific) for 5 min, followed by excessive rinsing with $H_2O$. Substrates were blown dry with air before attaching to the flow chamber system. SLBs were formed on substrates by incubating the prepared SUVs at 0.25 mg/mL mixed in 0.5x tris buffered saline (TBS) (1x TBS: 20 mM Tris, 136 mM NaCl, pH 7.4) for at least 30 min. Chambers were then rinsed with 1x TBS buffers. Next, 1 mg/mL (0.1%) BSA (Sigma) in TBS buffers was incubated for 10 min to block detects in SLBs.

Then solutions were exchanged to borate buffered saline (BBS) buffers (10 mM sodium borate, 150 mM NaCl, pH 8.5) (Rockland). Reduced DNAs (see *DNA Preparation*) were then added to the solution at a desired concentration (~50-200 nM for a typical titration experiment) and incubated for 1 hr (higher densities were achieved with higher concentrations and up to 2.5 hr of incubation). The sample was then heavily washed in the last step with PBS buffers (the imaging condition). All bilayer preparation was done at room temperature, unless stated otherwise.

Fluorescence Correlation Spectroscopy

FCS data were collected with an inverted Zeiss LSM 780 multiphoton laser scanning confocal microscope. Samples were excited by a 488 nm laser line (3 μW) that passed through a 488 beam splitter and focused by a C-APO 40x (FCS-certified) water-based objective. Emissions passed through a variable secondary dichroic (VSD) beam splitter that selected signals from wavelength 499-579 nm. The pinhole was aligned using the Adjust Pinhole function in the software. Signals were collected by the LSM BiG module with GaAsP photodetector, which enabled the selection of detection wavelength. All data were acquired using the ZEN Black software.

The ZEN program calculated time autocorrelation functions by $G_{ZEN}(\tau) = \frac{\langle I(t)I(t+\tau)\rangle}{\langle I(t)\rangle^2}$, where $I(t)$ is the fluorescence intensity, $\tau$ is the delay time, and $\langle\cdot\rangle$ denotes the average. This function relates to our definition of autocorrelation function by $G(\tau) = \frac{\langle \delta I(t)\delta I(t+\tau)\rangle}{\langle I(t)\rangle^2} = G_{ZEN}(\tau) - 1$, where $\delta I(t) = I(t) - \langle I(t)\rangle$ (*4*). Subsequent fitting was performed in Igor Pro (version 6). A Brownian diffusion model was used to fit the autocorrelation data:

$$G_{3D}(\tau) = \frac{1}{N\left(1+\frac{\tau}{\tau_D}\right)\sqrt{\left(1+\frac{1}{s^2}\frac{\tau}{\tau_D}\right)}} \quad \text{Eq. S3}$$

where $N$ is the particle number, $\tau_D$ is the characteristic diffusion time, and $s$ is the structural parameter of the optics and was fixed to 7 as suggested by the manual. For membrane experiments, autocorrelation functions were fitted by a 2D diffusion model: $G_{2D}(\tau) = \frac{1}{N(1+\tau/\tau_D)}$. All curve fitting was performed between a time range of 10 μs to 1 s (except spot-size calibrations, where a time range of 1 μs-0.1 s was used). The diffusion coefficient of the sample was calculated by $D = w^2/4\tau_D$, where $w$ is the confocal spot radius and $\tau_D$ is the measured diffusion time. The spot-size ($w$) was calibrated by averaging spot-sizes determined by fluorescein (Sigma Aldrich), Alexa Fluor 488 (Invitrogen), Atto488-carboxylic acid (Atto-Tec), and 0.1 μm TetraSpeck microspheres (Invitrogen) in water at room temperature assuming known diffusion coefficients ($D = 425, 435, 400,$ and 4.4 μm$^2$/s, respectively). For measurements on supported membranes, it is worth noting that reliable autocorrelation functions and stable intensities are strong indicators of fluid membranes (Fig. S3). We did not see any discernable photobleaching in the intensity traces (even when moving to a new position), suggesting that DNAs were mostly fluid on bilayers. All DNA densities on SLBs were measured independently prior to strand-displacement reactions.

Epifluorescence Microscopy

Monitoring strand-displacement reactions on supported membranes was performed on a Nikon TiE inverted microscope. Samples were excited by a 475/28-nm light source (Lumencor Spectra III) (120 uW). The excitation light passed through a 474/27-nm bandpass filter and a 493-nm dichroic mirror (Sedat quad filter set, Semrock) and focused through a 100x 1.49 N.A. oil immersion objective (Nikon). Emission signals passed through a 528/38-nm bandpass filter (Sedat quad filter set, Semrock) and were collected by an Andor EM-CCD camera (iXon DU-897). Images were acquired every 10 s with an exposure time of 200 ms. All data acquisitions were controlled by the NIS-Elements software. Prior to imaging, the sample was buffer exchanged to PBS buffers containing 10 mM 2-Mercaptoethanol (BME) (Sigma) and 2 mM Trolox (Cayman Chemical) to reduce photobleaching. Strand-displacement reactions were triggered by including initiator strands (I) in the above PBS buffers. Photobleaching was estimated with samples without the initiator strand. Fluorescence intensities from images were extracted in ImageJ. After background subtraction, the intensities were normalized to the initial point. Data were then corrected for photobleaching by dividing each time point by the photobleaching curve. Processed data were then fitted by the same procedure described in the *Fluorometer* section (Eq. S2).

Unit and Unit Conversion

The rate constants $k_i$ derived from solution and membrane experiments were expressed in the unit of $M^{-1}s^{-1}$ and $\mu m^2 s^{-1}$, respectively. The solution rate constants ($k_s$, $k_F$, and $k_c$) were then converted to the unit of $\mu m^3 s^{-1}$ by $M^{-1} = \frac{dm^3}{mole} = \frac{10^{15}\ \mu m^3}{6\times 10^{23}\ molecule} = 1.67 \times 10^{-9} \frac{\mu m^3}{molecule}$. Thus, $k_s = 2.7 \times 10^5 M^{-1}s^{-1} = 4.5 \times 10^{-4}\ \mu m^3 s^{-1}$; $k_F = 1.6 \times 10^5 M^{-1}s^{-1} = 2.7 \times 10^{-4}\ \mu m^3 s^{-1}$; $k_c = 1.3 \times 10^4 M^{-1}s^{-1} = 2.2 \times 10^{-5}\ \mu m^3 s^{-1}$. In all subsequent analyses, rate constants have units in µm and s.

Derivation of distance to the nearest target

In the following, we derive the distance distribution of a single signaling molecule with its nearest target molecule. Particles are randomly distributed and non-interacting. Following the seminal work of S. Chandresekhar (*5*), we define $w_{3D}(x)$ as the probability of finding the nearest neighbor molecule between $x$ and $x + dx$ in the solution, with the fluorescent molecule marked at the center. By definition:

$$w_{3D}(x) = \left(1 - \int_0^r w_{3D}(x')dx'\right)4\pi x^2 c. \qquad \text{Eq. S4}$$

Note that the concentration adopts a unit of $\mu m^{-3}$. Eq. S4 is solved by differentiation and then integration with respect to $x$:

$$\frac{d}{dx}\frac{w_{3D}(x)}{4\pi x^2 c} = \frac{d}{dx}\left(1 - \int_0^x w_{3D}(x')dx'\right) = -w_{3D}(x) = -4\pi x^2 c \left(\frac{w_{3D}(x)}{4\pi x^2 c}\right),$$

$$\int \frac{d\left(\frac{w_{3D}(x)}{4\pi x^2 c}\right)}{\left(\frac{w_{3D}(x)}{4\pi x^2 c}\right)} = \int -4\pi x^2 c dx,$$

$$ln\left(\frac{w_{3D}(x)}{4\pi x^2 c}\right) = -\frac{4}{3}\pi x^3 c, \qquad \text{Eq. S5}$$

from which we obtain:

$$w_{3D}(x) = 4\pi x^2 c e^{-4\pi x^3 c/3}. \qquad \text{Eq. S6}$$

$w_{3D}(x)$ describes the full statistical distribution of the nearest target (Fig. 4D). Thus, the average distance is:

$$d = \langle x \rangle = \int_0^\infty r w_{3D}(r) dr = \Gamma\left(\frac{4}{3}\right)\left(\frac{3}{4\pi c}\right)^{1/3} \approx 0.554\mathrm{c}^{-1/3},$$

$$c = \frac{0.17}{d^3}, \qquad \text{Eq. S7}$$

where Γ is the gamma function.

The nearest-neighbor distribution in 2D is defined similarly:

$$w_{2D}(x) = \left(1 - \int_0^x w_{2D}(x')dx'\right) 2\pi x\sigma, \qquad \text{Eq. S8}$$

which can be solved following steps in Eq. S5:

$$\frac{d}{dr}\left(\frac{w_{2D}(x)}{2\pi x\sigma}\right) = \frac{d}{dx}\left(1 - \int_0^x w_{2D}(x')dx'\right) = -w_{2D}(x) = -\left(\frac{w_{2D}(x)}{2\pi x\sigma}\right) 2\pi x\sigma,$$

$$\int \frac{d\left(\frac{w_{2D}(x)}{2\pi x\sigma}\right)}{\left(\frac{w_{2D}(x)}{2\pi x\sigma}\right)} = \int -2\pi x\sigma dx,$$

$$\ln\left(\frac{w_{2D}(x)}{2\pi x\sigma}\right) = -\pi x^2\sigma, \qquad \text{Eq. S9}$$

from which we arrive: $w_{2D}(x) = 2\pi x\sigma e^{-\pi x^2\sigma}$. Eq. S10

Thus, the average distance in 2D is:

$$d = \langle x \rangle = \int_0^\infty x w(x) dx = \Gamma\left(\frac{3}{2}\right)\left(\frac{1}{\pi\sigma}\right)^{1/2} = 0.5\sigma^{-1/2},$$

$$\sigma = \frac{1}{4d^2}, \qquad \text{Eq. S11}$$

At this point, it is useful to calculate some numbers relevant to cells. An average protein in cells has a concentration of about 200 nM (assuming 1/25,000 of total proteins), which is equivalent to an average distance of about 112 nm; a distance of 100 nm maps to a surface density of about 25 molecules·$\mu m^{-2}$.

### Supplementary Figures

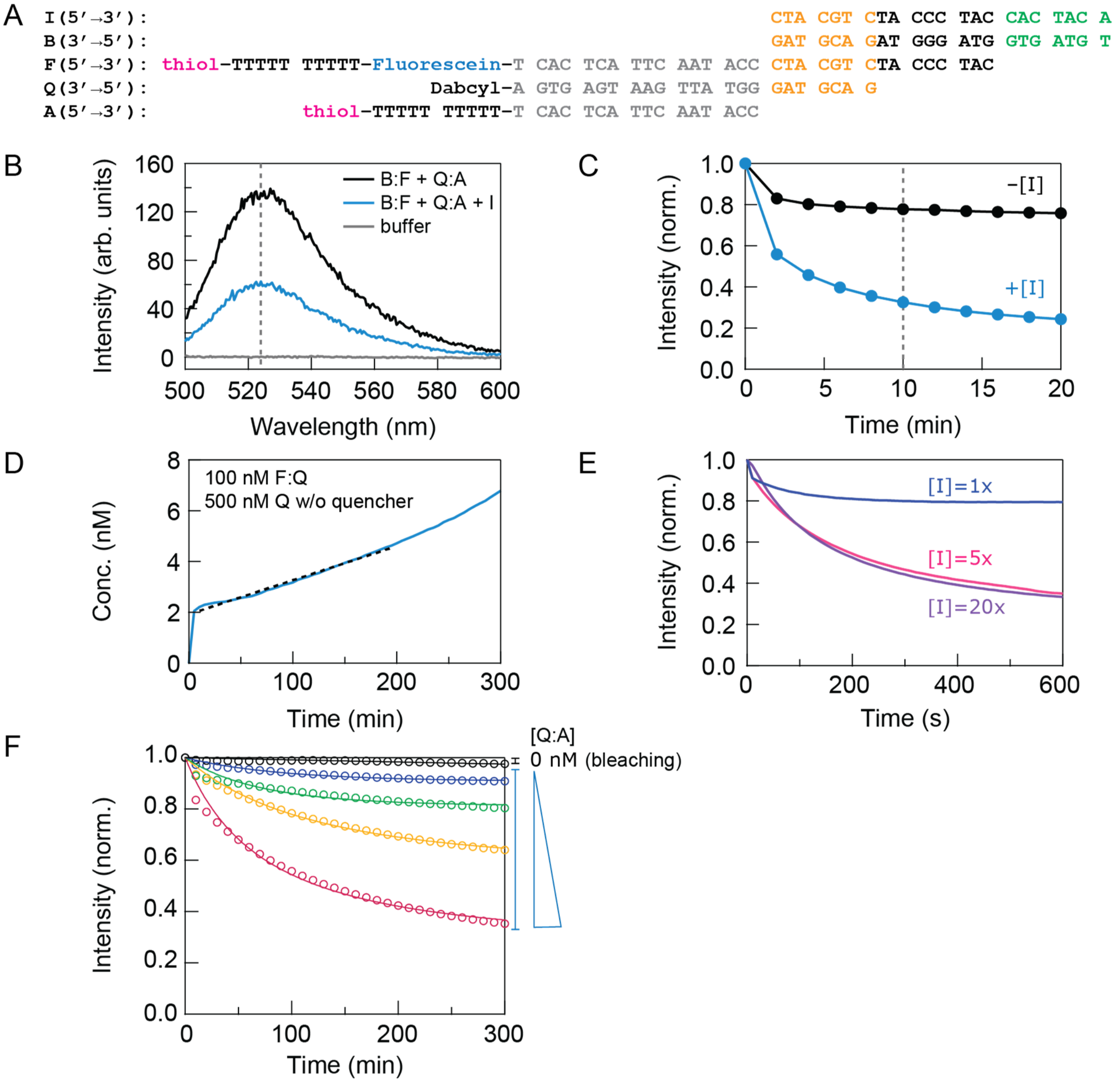


**Fig. S1. Characterization of strand-displacement reactions in solution.**

(**A**) DNA constructs used in this study. The fluorescein derivative used was FAM. When coupled to the membrane, the left side of the sequence was closest to the membrane surface (the thiol group is the coupling site). (**B, C**) Fluorescence spectra and time courses of the blocked fluorescent strand (B:F) with and without the initiator strand (I). Dashed lines show the wavelength used to measure the time course (left) and the time point when the spectrum was taken (right). (**D**) Estimation of the upper bound of the dissociation rate of DNA complexes (F:Q). To resolve the dissociated strands, we added complementary strands without the quencher such that most strands, once dissociated, will remain fluorescent. By fitting a linear rate to the early slope (slope $\approx k_{-1}[F:Q]$), the fit yielded a dissociation rate constant of $2 \times 10^{-6}$ $s^{-1}$. Note that this estimation is likely higher than the true dissociation rate because the additional complementary strand may compete with the quencher strand (Q). Nevertheless, this slow rate indicates that the average lifetime of these DNA complexes ($\lesssim 10^6$ s) are much longer than the time course of titration experiments ($\lesssim 10^3$ s) (Fig. 2). Therefore, in these experiments, the number of association events can be approximated by the number of complexes. (**E**) Titration of the initiator strand (I) on the kinetics of strand-displacement reactions. When the initiator's concentration was above 5-fold of the fluorescent strand (F), the kinetics became mostly independent of the initiator's concentration, which suggests that the rate-limiting step was the fluorescent strand (F) binding to the quencher strand (Q). (**F**) Data in Fig. 2B overlaid with photobleaching control (black) under the same experimental condition. At the end of the experiment, less than 5% of molecules were photobleached.

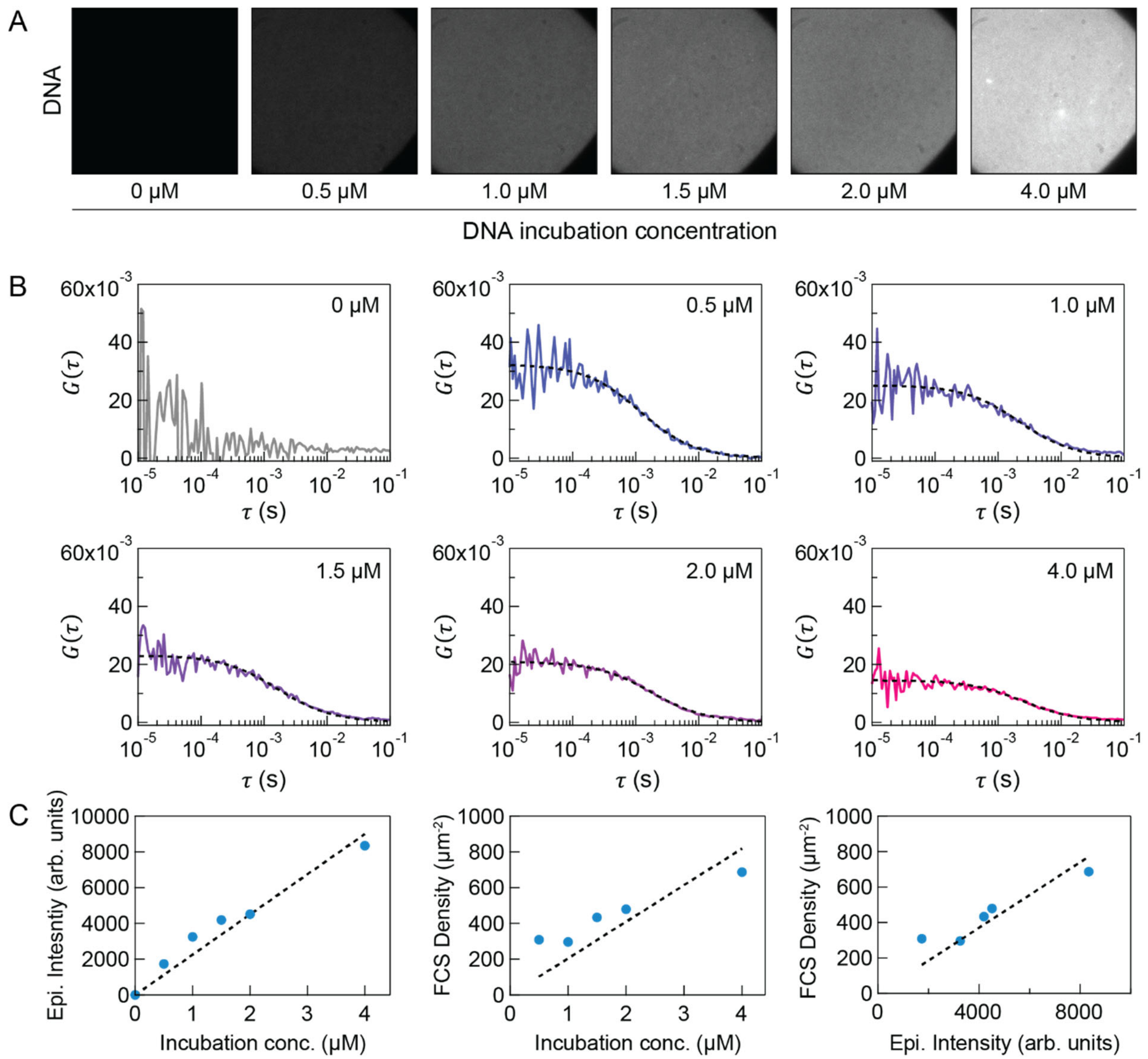


**Fig. S2. DNA density titration on supported membranes**

(**A**) Epifluorescence images of titrating the fluorescent strand (F) on supported membranes. (**B**) Surface densities of DNA were calibrated by fluorescence correlation spectroscopy (FCS). Dashed lines are fitting of a 2D Brownian model. (**C**) Summary of data shown in (**A**) and (**B**). It was experimentally convenient to have a linear relationship (dashed lines) between surface densities and incubation concentrations, though this linearity was not required for our experiments. We measured the density of all the samples independently immediately prior to strand-displacement reactions.

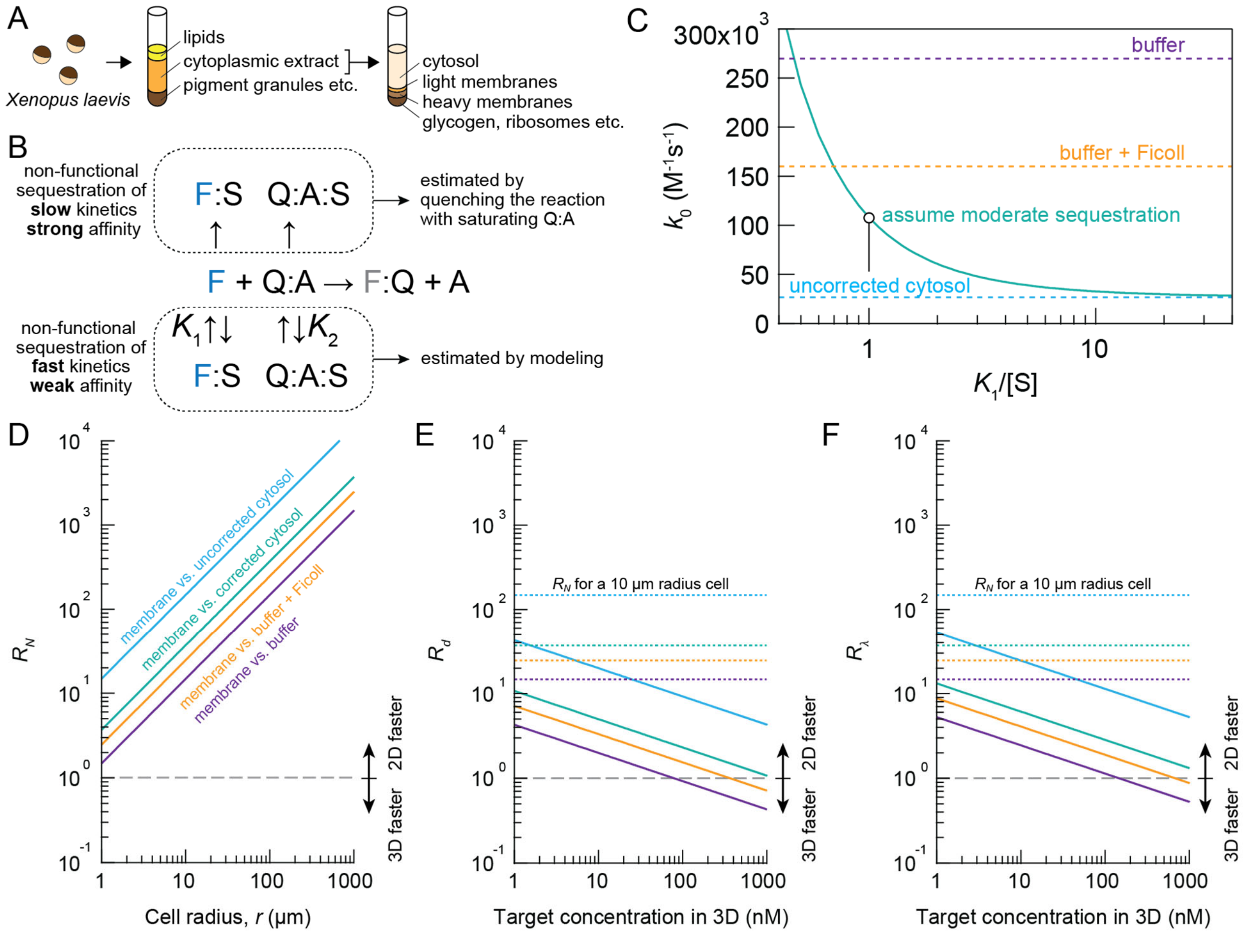


**Fig. S3. Association measurements and analysis in cytosolic extracts.**

(**A**) Schematic of cytosolic extract protocols. *Xenopus laevis* eggs were harvested, crushed, and fractionated using a centrifuge to obtain cytoplasmic extracts (the middle layer of the middle tube), which contain cytosolic proteins and membrane organelles. By further fractionating cytoplasmic extracts using ultracentrifugation, cytosolic proteins (top layer) were separated from membrane organelles. Cytosolic extracts were taken and frozen down for future use. Note that cytosolic extracts prepared as such are essentially undiluted. (**B**) Sequestration of DNA probes in cytosolic extracts. The functional fraction of DNA probes, defined by the ability to perform strand-displacement reactions, in extracts was about 60-70%. This functional fraction was estimated by measuring fluorescent intensities at equilibrium with excess quencher strands (Q:A). Data were normalized by this functional fraction, which presumably correct for any sequestration of high affinity with slow kinetics. Given the measured slow rate compared to crowding agent solution, we suspect that additional sequestration with fast kinetics, possibly in the form of nonspecific binding, are present in extracts, which lowers the concentration of free DNA. We modeled this effect by assuming this nonspecific sequestration to be in quasi-equilibrium with the free DNA probes. The rate equation is $\frac{dx}{dt} = k_0(F_{tot} - x)\left(\frac{K_1/[S]}{1+K_1/[S]}\right) * (Q_{tot} - x)\left(\frac{K_2/[S]}{1+K_2/[S]}\right)$ where $k_0$ is the true association rate constant, $K_i$ is the dissociation constant, and $[S]$ is the concentration of sequestration molecules. Assuming that $[S]$ is in excess to the DNA, solving this model gives an identical solution to a bimolecular reaction, except the rate constant is modified by $k =$

$k_0\left(\frac{K_1/[S]}{1+K_1/[S]}\right)\left(\frac{K_2/[S]}{1+K_2/[S]}\right)$. (**C**) Correction for sequestration of fast kinetics, using the model from (**B**) and assuming that $K_1 = K_2$. If we consider the case of moderate sequestration, $K_1/[S] = 1$ (50% sequestrated) (circle), the corrected rate constant is $1.1\times10^5$ $M^{-1}s^{-1}$, or $1.8\times10^{-4}$ $\mu m^3 s^{-1}$, which is close to the values in crowding agent solutions ($1.6\times10^5$ $M^{-1}s^{-1}$). (D-F) Estimates of R metrics with and without correction in cytosolic extracts. R metrics are defined in the main text, also see Fig. 4.

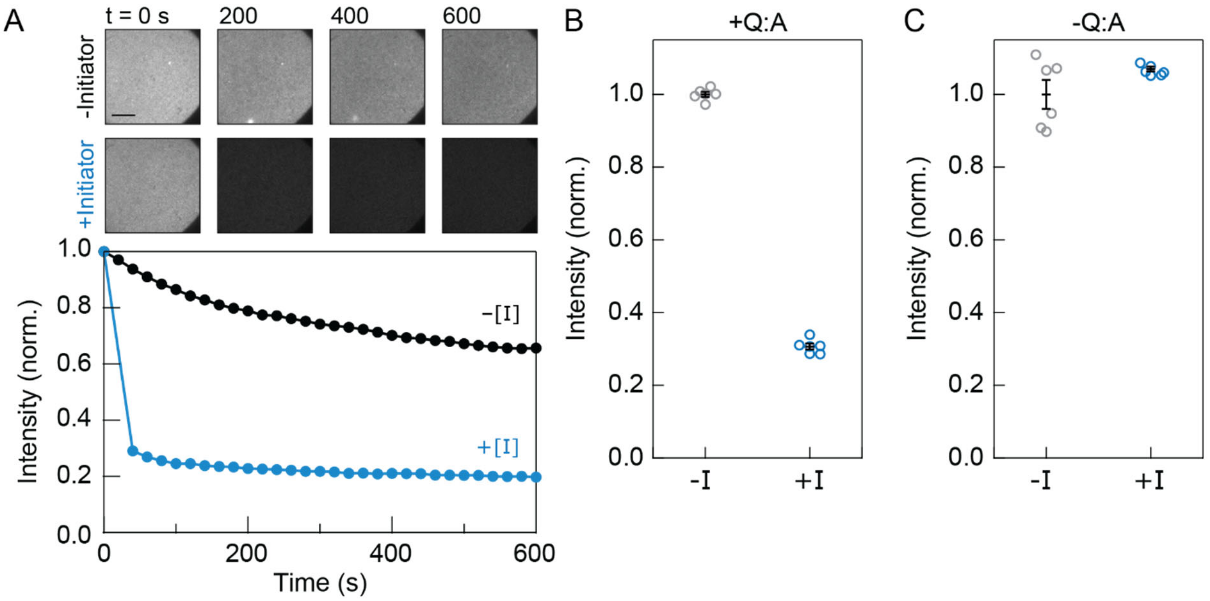


**Fig. S4. Characterization of strand-displacement reactions on supported membranes.**
(**A**) Images (top) and time courses (bottom) of strand-displacement reactions with and without the initiator strand (I) on supported membranes. Note that the initiator-dependent fluorescent decay (blue) was much steeper than the bleaching curve (black) (prior to photobleaching correction). Scale bar, 20 µm. (**B**) Epifluorescence intensities before and after adding initiator at different regions of interest. In contrast to (**A**), intensities here were measured without prior exposure to light (i.e., no photobleaching). (**C**) Same as in (**B**), except without the quencher strand (Q). Error bars, SEM from 6 different region of interests (ROI).

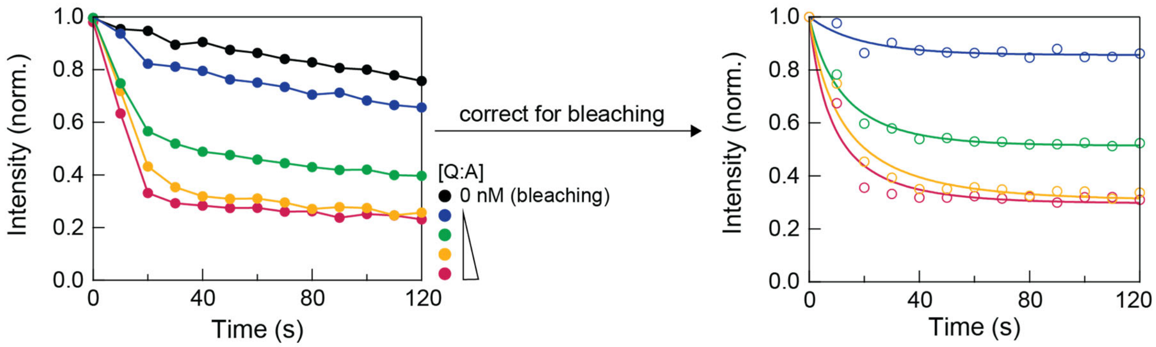


**Fig. S5. Photobleaching correction for reactions on supported membranes.**

Normalized intensity traces before (left) and after (right) photobleaching correction. Data were corrected by dividing the intensity of each time point by that of the bleaching curve (black). Note that, after correction, the kinetic traces plateaued near the end of experiments, suggesting that the correction was appropriate.

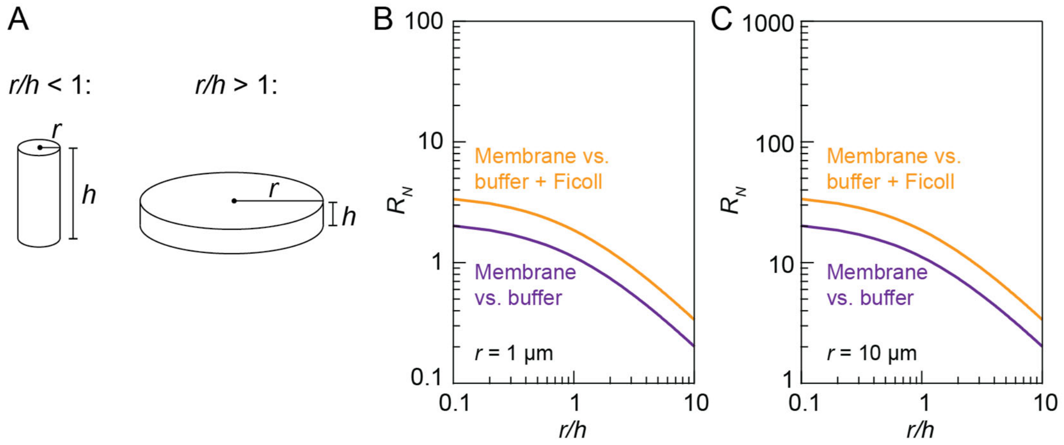


**Fig. S6. Other geometry for $R_N$: cylinders and disks.**

(**A**) Geometry of a cylinder and a disk. Both shapes are parameterized by radius and height, but the difference is that cylinders have $r < h$, whereas disks have $r > h$. In both cases, $\frac{V}{A} = \frac{rh}{2(r+h)}$ and $R_N = \frac{k_{2D}}{k_{3D}} \frac{r}{3} \left( \frac{3}{2\left(1+\frac{r}{h}\right)} \right)$. (**B**, **C**) $R_N$ of cylinders and disks. For example, a bacterium of $r = 0.5$ μm and $h = 3$ μm gives $R_N \approx 1.6$; a mammalian cell of $r = 10$ μm and $h = 5$ μm gives $R_N \approx 12$.
